# Increasing proteome coverage using cysteine-specific DIA Mass spectrometry – Cys-DIA

**DOI:** 10.1101/2020.02.27.966861

**Authors:** Muhammad Tahir, Arkadiusz Nawrocki, Henrik J. Ditzel, Martin R. Larsen

## Abstract

We report a Data Independent Analysis (DIA) mass spectrometric method for quantitative proteomics based on cysteine-containing peptides derived by tryptic digestion from biological samples (termed Cys-DIA). This technology significantly increases the coverage of the identifiable and quantifiable proteome in DIA/SWATH analyses by >35% compared to the conventional DIA/SWATH methods. The strategy was applied here to illustrate the quantitative difference in the proteome between isogenic human cancer cell lines with different metastatic capabilities.

---

Technological advances in mass spectrometry-based proteomics still face a key challenge to maximize the coverage of the proteome to reproducibly detect low abundant proteins. Data independent acquisition (DIA/SWATH) is an unbiased technology in quantitative proteomics to monitor the entire detectable proteome from a complex biological sample using advanced liquid chromatography tandem mass spectrometric (LC-MSMS) instrumentation^1^. Peptides can be analyzed by LC-MSMS in DIA mode where m/z windows are selected (typical 25-50 windows covering m/z 400-1500) and all peptides within each window are fragmented, resulting in very complex fragmentation spectra. Peptides can then be identified and quantified using an external experimentally generated peptide spectral reference library (SRL) for searching. The SRL is produced by more traditional data-dependent acquisition (DDA or shot-gun) approaches and includes specific information on unique peptides derived from tryptic digestion of proteins including their HPLC retention time, peptide and fragment masses, used to interrogate DIA data. Depending on LC-MSMS instrumentation, gradient length and LC setup, the proteome coverage by DIA analyses is on average from 2,500-5,000 proteins for eukaryote cells, tissues and mixed cell populations. New strategies that increase the proteome coverage in DIA would make the method more attractive to applications in the biomedical field.

Here we describe a new DIA method in which the DIA is performed using only cysteine-containing peptides (Cys-peptides). Cysteine is the second rarest amino acid, but is present in about 97% of human proteins^2^. In conventional shot-gun proteomics, cysteines are chemically modified by reduction and then alkylation with iodoacetamide to make stable uniform Cys-peptides that are used for identification and quantitation equally with other unmodified peptides. Due to the ability to chemically manipulate Cys residues, several techniques have been reported capitalizing on this property. One of the first isotope labeling strategies in proteomics used a chemical tag directed against Cys residues, the isotope-coded affinity tags (ICAT)^3^. Pairs of cysteine-binding ICAT tags differ in molecular mass and are used to differentially label two samples of proteins or peptides. Labeled peptides can then be purified and analyzed by LC-MSMS. This allows accurate quantitation of sample pairs by targeting only the Cys-peptides for quantitation.

The Cys-DIA strategy provide higher coverage of the proteome. The most unique aspect of the new strategy is the production of a Cys-peptide specific SRL (Cys-pep-SRL) to search the DIA/SWATH data obtained from purified Cys-peptides. The Cys-pep-SRL is generated by purifying Cys-peptides using Thiol disulfide exchange (TDE) chromatography^4^ (**Fig. 1A**) followed by fractionation of the Cys-peptides after alkylation with iodoacetamide using high pH reversed-phase fractionation (HpH RP) into 20 concatenated fractions^5^, that are then analyzed by LC-MSMS using DDA. Here, sample specific Cys-pep-SRLs were generated from tryptic digestions of HeLa cells and a mixture of isogenic cancer cell lines (NM2C5, M4A4 and CL16) using 60 min gradients (EASY-LC) and 21 min gradients (EASY-LC and EvoSep-LC), respectively. For comparison with traditional DIA a tryptic peptide SRL from HeLa cells were obtained from Thermo Fisher Scientific (Whole-SRL). The Whole-SRL was made using the same 60 min gradient (EASY-LC) as in this study.

**Figure 1.**
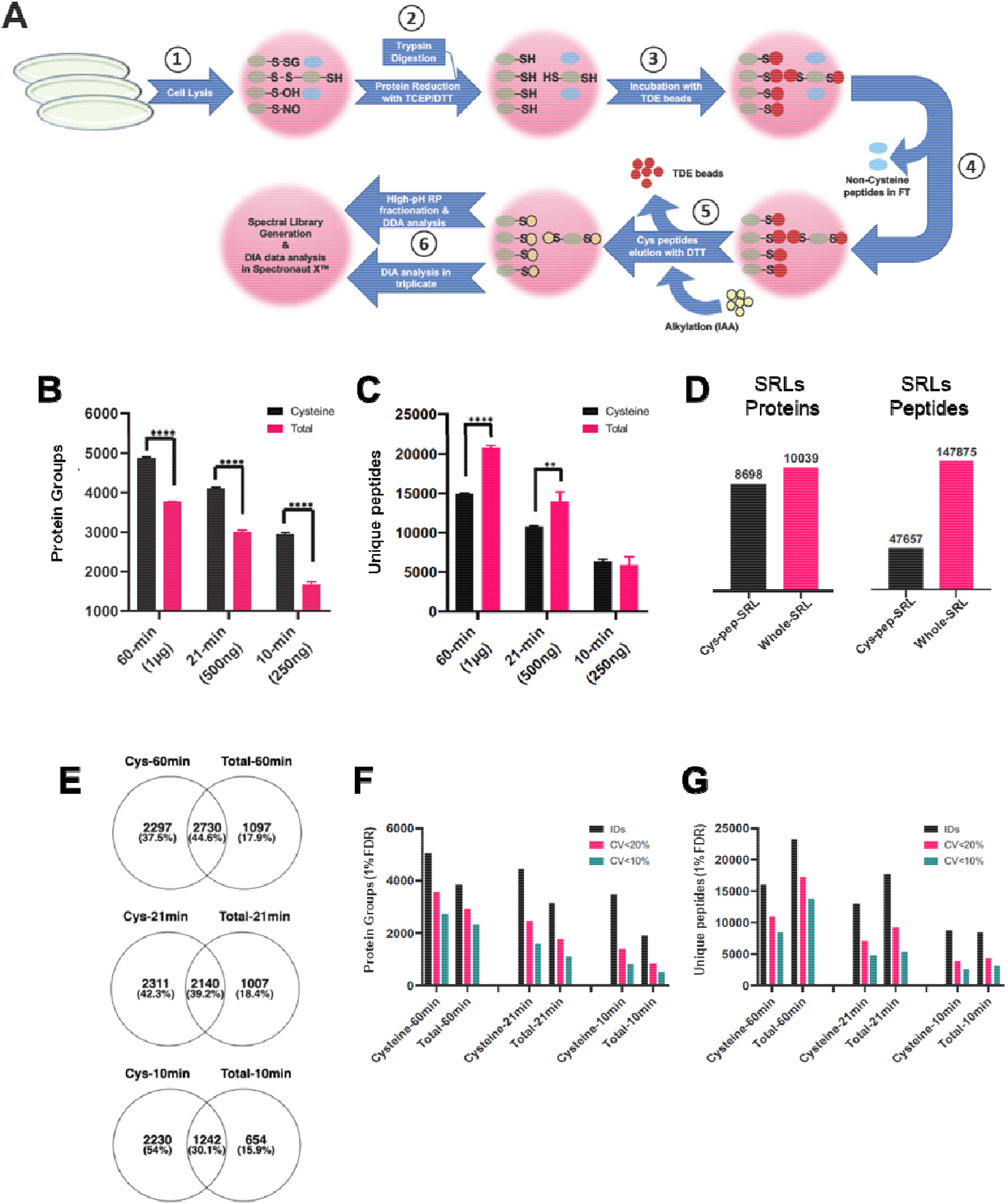
**(A)** A schematic presentation of the workflow for sample processing for enrichment of Cys-peptides. **(B)** Number of identified protein groups in HeLa cell tryptic peptide mixtures using Cys-DIA (Cysteine) and normal DIA (total). The identifications were made from triplicates using 60min, 21min and 10min of capillary chromatographic separation coupled with HRMS1 DIA on a Q-Exactive HF-X mass spectrometer. **(C)** Number of identified unique peptides in the above-mentioned experiments. **(D)** Number of proteins and peptides present in the Cysteine and total spectral libraries. **(E)** Overlap between Cys-DIA and total DIA for the three chromatographic separation times. **(F)** Number of identified and quantified (<20% CVs and <10% CVs) proteins. **(G)** Number of identified and quantified (<20% CVs and <10% CVs) unique peptides.

The HeLa 60 min gradient Cys-pep-SRL contained 47,657 unique Cys-peptides, covering 8,698 proteins (**Fig. 1D**), whereas the EvoSep 21 min cancer cell line Cys-pep-SRL contained 7073 unique Cys-peptides covering 3887 proteins. The HeLa whole-SRL contained 147,875 unique peptides from 10,039 proteins (**Fig. 1D**).

The performance of the Cys-DIA strategy was investigated using purified Cys-peptides and tryptic peptides from HeLa cells (1µg peptide solution applied in both cases), using High Resolution MS1 DIA (HRMS1-DIA), where the peptide quantitation is performed on the MS level. All analyses were performed with three technical replicates. The Cys-DIA method, searched against the Cys-pep-SRL, allowed for the identification of 4,869 ± 15 proteins, whereas the standard 60 min DIA, searched against the Whole-SRL, resulted in the identification of 3,767 ± 7 proteins **(Fig. 1B**). This corresponds to an increase in the proteome coverage of 30%. By applying a shorter 21 min gradient (EASY-LC) and 500 ng material, using the 60 min SRLs and retention time normalization peptides^6^, we identified 4,097 ± 36 using Cys-DIA, but only 2,997 ± 39 proteins using standard DIA (**Fig. 1B**). The increase in the proteome coverage was 36.9%. Decreasing the gradient to 10 min (EASY-LC) and the peptide loading to 250 ng resulted in an increase in the proteome coverage of 76% (2,928 vs 1,663 proteins, 54% new proteins in the Cys-DIA (**Fig. 1E**)) by using the Cys-DIA method and the 60 min Cys-pep-SRL with retention time normalization peptides (**Fig. 1B**).

The enrichment of Cys-peptides significantly reduces the complexity of the sample prior to Cys-DIA by eliminating the non-Cys peptides and the Cys-DIA method resulted in less identified unique peptides for both the 60 min and 21 min gradients compared with the standard whole-SRL (14,887 vs. 20,597 and 10,689 vs. 13,960, respectively). The 10 min gradient had equal amounts of unique peptides identified in Cys-DIA and standard DIA (**Fig. 1C**). The latter is expected, as the Cys-peptide mixture from a complex sample is still highly complex.

The overlap between all proteins identified from the technical triplicate study at the protein level was 44.5 %, 39,2% and 30,1% for the 60 min, 21 min and 10 min gradient, respectively **(Fig. 1E)**. From these data it is observed, that if the same HeLa sample was analyzed by using the normal DIA (Whole-SRL) and Cys-DIA (Cys-pep-SRL) a total of 6,124 proteins could be identified compared to only 5,027 proteins from the sum of the technical triplicate analysis using normal DIA alone.

The number of quantifiable proteins based on the Cys-peptides also increased relative to the standard DIA. The increase in the number of quantifiable proteins with 10% CVs was for the 60 min, 21 min and 10 min gradients 16.5%, 43.3% and 58.7%, respectively (**Fig. 1F**). This illustrates the benefit of using the Cys-DIA method for quantitative proteomics.

The Cys-DIA method was applied to determine the quantitative difference in the proteomes of an isogenic metastasis model of human cancer, in which the 3 different isogenic cell lines exhibited different metastatic capabilities^7, 8^. A new sample-specific peptide SRL (Cys-cancer-SRL) was generated from 20 concatenated high pH RP fractions of Cys-peptides purified from a tryptic peptide mixture derived from proteins from a mixture of the three cancer cell lines; non-metastatic (NM2C5), weakly metastatic (M4A4) and highly metastatic (CL16). The 21 min Cys-cancer-SRL using an EvoSep LC system consisted of 7073 unique Cys-peptides covering 3887 proteins.

Subsequent 21 min Cys-DIA of purified Cys-peptides from each of the cancer cell lines (biological triplicates) using an EvoSep LC system resulted in the identification of about 2,700 protein groups from each cell line, of which about 2,000 were quantified across all replicates with an average CV of 11 % (**Fig. 2A**). On average about 4,500 Cys-peptides were identified in the triplicate analysis of the 3 different cell lines with an average CV of 11.5 % (**Fig. 2B**).

**Figure 2.**
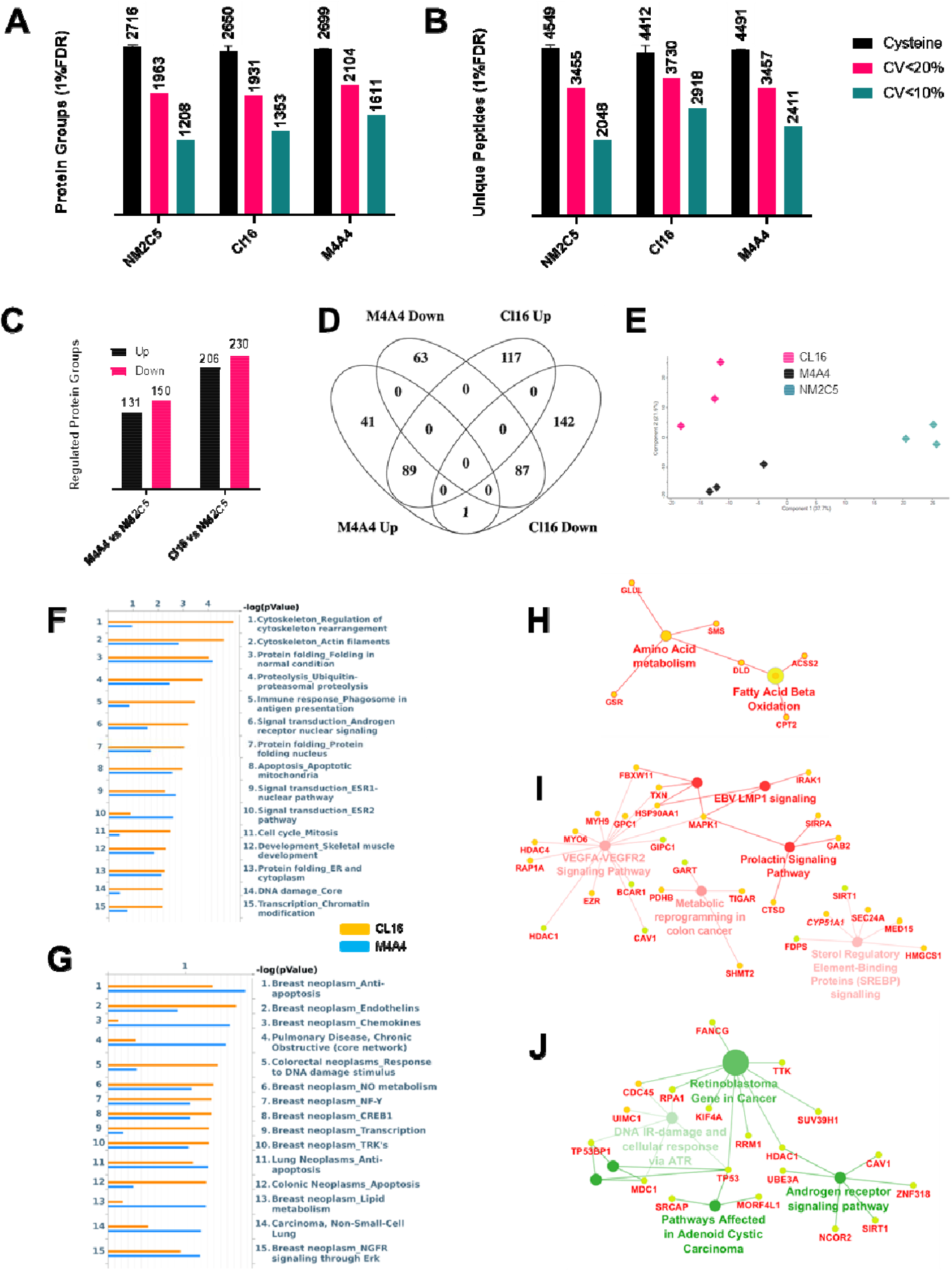
**(A)** Number of identified protein groups in the three cancer cell lines using Cys-DI including the number of quantifiable proteins with CVs <20% and <10%. **(B)** Number of identified and quantified unique peptides in the experiment. **(C)** Number of regulated proteins in the M4A4 and CL16 cell lines in comparison to the lowest metastatic cell line NM2C5. The regulated peptides were selected with a *p*-value less than 0.01 and a fold change cutoff of 2. **(D)** Overlap between up- and down-regulated proteins in M4A4 and CL16 in comparison to NM2C5. **(E)** PCA plot based on the identified and quantified proteins in the three cancer cell lines. Sea color represents the NM2C5 cell line, black represents the M4A4 cell line and pink represents the CL16 cell line. **(F)** Enriched GO annotation pathway analysis for regulated proteins in M4A4 and CL16. **(G)** Disease biomarker networks analysis for regulated proteins in M4A4 and CL16. **(H)** Example of a highly upregulated pathway in M4A4 compared with NM2C5. **(I)** Example of highly upregulated pathways in CL16 compared to NM2C5. **(J)** Example of highly downregulated pathways in CL16 compared to NM2C5. WikiPathways analysis was performed by using ClueGO/CluePedia as a Cytoscape plug-ins.

The list of total cysteine-containing proteins was subjected to statistical test using the ShinyApp^9^. A threshold of *p*-value less than 0.01 and a fold change cutoff of 2 were used for significant regulation of proteins. The analysis resulted in 281 and 436 significantly regulated proteins in M4A4 and CL16 compared to the non-metastatic NM2C5 cell line, respectively (**Fig. 2C**). An overlap for the common and unique significantly regulated proteins was observed in the different cancer cell lines (**Fig. 2D**). The PCA plot showed that the replicates are grouped and separated according to cell type (**Fig. 2E**).

The regulated proteins in CL16 versus NM2C5 were compared to regulated proteins identified in a previous study with significant lower proteome coverage (414 proteins)^8^. Only five regulated proteins were found to be overlapping between the two data sets, however, all were regulated in the same direction, validating the Cys-DIA method for quantitative proteomics.

Analysis of the regulated proteins in CL16 and M4A4 in comparison to the NM2C5 showed increased regulation of pathways associated with cytoskeleton, protein folding, proteolysis, cell cycle, and apoptosis (**Fig. 2F**), correlating well with disease biomarker networks (MetaCore) related with breast cancer development and metastasis (**Fig. 2G**).

Pathway analysis using the Wiki-pathway identified amino acid metabolism and fatty acid beta oxidation as highly upregulated in M4A4 compared with NM2C5 (**Fig. 2H**), reflecting the increasing metabolic demand in these cells^10, 11^. Vascular endothelial growth factor (VEGF), latent membrane protein 1 (LMP1) and prolactin signaling pathways were significantly upregulated in CL16 compared to NM2C5 (**Fig. 2I**). These pathways are important for cancer growth, development and angiogenesis, characteristic for a cell at a high metastatic level.

In the pathways of highly downregulated proteins in CL16 compared to NM2C5 we identified tumor suppressor p53 (TP53) and TP53-binding protein 1 (TP53BP1) (**Fig. 2J**), which are involved in a number of processes such as induction of growth arrest, DNA repair and apoptosis^12^, correlating well with the metastatic level of this cell line.

In conclusion, we have shown that the new Cys-DIA strategy, wherein the purified Cys-peptides are used for DIA mass spectrometry identification and quantitation significantly increase the proteome coverage compared with conventional DIA/SWATH analysis where all peptides are used. The increase in the proteome coverage is particularly significant when using shorter LC gradients, as illustrated with both EvoSep/Easy LC 21 min/10 min LC gradients, respectively. This method will be very useful in quantitative DIA proteomics studies where large number of samples need to be analyzed using DIA with shorter gradients, such as in clinical proteomics and personalized medicine.

## Materials & Reagents

All chemicals were purchased from Sigma-Aldrich (St. Louis, MO), unless otherwise stated. Trypsin from porcine pancreas was from Sigma and modified in house with dimethyl labeling. Poros Oligo R3 reversed-phase material was from Applied Biosystems (Forster city, CA). The 3M Empore™ C8 disk was purchased from 3M Bioanalytical Technologies (St. Paul, MN). OASIS HLB columns were purchased from Waters. Thiopropyl Sepharose™ 6B beads were from GE Healthcare. Easy spray column (ES806, 15 cm & ES802A, 25 cm) were from Thermo Fischer Scientific. All solutions were made with ultrapure Milli-Q water (Millipore, Bedford, MA).

### Samples

Human cervix epithelial adenocarcinoma (HeLa) cells and isogenic cancer cell lines (NM2C5, M4A4 and CL16) were cultured as described previously^13, 14^.

### Protein Extraction

Proteins were extracted from HeLa and cancer cells by using 6M Urea and 2M thiourea lysis buffer containing the 10mM DDT, and protease inhibitors cocktail. The cells were lysed using 5 cycles of tip sonication with 40% output. Each cycle was for 15 seconds with 1 minutes of interval on ice. After sonication, the lysate was kept for about 1 hour at room temperature to maximize the protein solubility. Protein quantification was performed using Qubit (Thermo Fisher Scientific).

### Reduction, trypsin digestion and peptide cleaning

The protein samples with 6M urea/2M thiourea were diluted 8 – 10 times with 20mM TEAB and trypsin was added 1:50 (trypsin:proteins). The samples were incubated at room temperature overnight. The tryptic peptides were purified using the commercially available OASIS HLB columns to remove the DTT. The bound peptides were eluted with 70% acetonitrile and the samples were dried down in a vacuum centrifuge.

### Cysteine enrichment and purification

The cysteine enrichment was performed as described previously^15^ with some modifications. A total of 30 mg of Thiopropyl Sepharose™ 6B beads were rehydrated in 1 ml of Milli-Q water twice, followed by 2 incubations with the 40% acetonitrile/20mM TEAB buffer, pH = 8.0. The dried reduced peptides were resuspended in the same buffer and incubated with the beads for 2 hours at room temperature with gentle agitation. After incubation, the flow-through was saved and the beads were washed 7 times with 40% acetonitrile/20mM TEAB buffer to remove unbound peptides. The Cys-peptides bound to the beads were eluted with 20mM DTT in 20 mM TEAB, pH 8. The eluted peptides were alkylated with 40mM iodoacetamide (IAA) in the dark, at room temperature. After alkylation, the peptide solution was acidified using 1% Formic acid and subsequently purified with homemade Poros Oligo R3 reversed-phase material microcolumns^16^ in p200 tips and dried down in a vacuum centrifuge prior to analysis.

### High pH reversed-phase fractionation

In order to generate peptide spectral libraries (SRLs), peptides (normal and Cys-peptides) from HeLa cells and cancer cell lines were dissolved in solvent A (20 mM ammonium formate, pH 9.2). Peptides were separated on an ACQUITY UPLC M-class, CSHTM C18, 130, 1.5m, 300 mm × 100 mm column (Waters), using a DIONEX Ultimate 3000 LC system (Thermo Fisher Scientific) equipped with a fraction collector. Peptides were separated in a linear gradient from 2 to 10% solvent B (2 volumes of solvent A added to 8 volumes of acetonitrile, pH not adjusted) in 5 min, then to 40% in 65 min, to 50% in 32 min, to 95% in 8 min, kept at 95% for 10 min and at 2% for 16 min for cysteine containing peptide fraction and total peptide fraction. A total of 52 fractions were concatenated into 20 for HeLa and cancer cell peptides. The fractions were dried down and resuspended in LC-MSMS solvent A (0.1% formic acid) supplemented with 1x concentrated iRT per injection (iRT Kit, BIOGNOSYS) prior to LC-MSMS. The Thermo HeLa SRL for the total proteome was provided by Thermo Fisher Scientific (Bremen, Germany) and it was generated from 36 concatenated high RP fractions on a Q-Exactive HFX using 1 hour gradients with iRT peptides included.

### Liquid Chromatography Tandem Mass spectrometry (LC-MSMS)

All LC-MSMS experiments were performed on a Q-Exactive HF-X mass spectrometer (Thermo Fisher Scientific (Bremen, Germany)) and the acquisition was either performed in DDA or DIA mode. For quality control and retention time normalization, reference peptides were spiked into all samples according to the manufacturer instructions (iRT Kit, Biognosis).

Chromatographic separations were performed using an EASY-nLC 1000 system (Thermo Fisher Scientific) and an Evosep LC system (Biosystems). Two types of biological samples (HeLa and cancer cell line) were used for analysis. The peptides were enriched for total Cys-peptides followed by high pH fractionation to generate sample specific SRLs.

Easy spray column of 150 ID was used from Thermo Fischer Scientific. The easy spray source temperature was set at 50°C. A 60, 21 and 10 min gradient (EASY-nLC) was used for HeLa and 21 min gradient (Evosep) for total cysteine of the cancer cell lines.

#### DDA runs

For peptide SRL generation, An EASY–LC system with either an EASY-SPRAY 25 cm, 75ID RP column (Thermo Fisher Scientific) or an EASY-SPRAY 15 cm, 150 ID (Thermo Fisher Scientific, Bremen, Germany) column, or an Evosep LC system (Evosep, Odense, Denmark) was used.

For the HeLa cell SRLs each High pH RP fraction was resolubilized in 0.1% Formic Acid and a fraction was loaded onto the EASY-SPRAY column. The peptides were eluted using a 60 min gradient using the EASY-LC system. The LC gradient used was: 2-8% B in 4 min, 8-32% B in 49 min, 32-60% B in 1 min at a flow rate of 300 nl/min (EASY SPRAY 75ID) or 1200 nl/min (EASY SPRAY 150 ID). The column was washed with 98% B. The column was heated to 50 °C. Mobile phases were as follows: (A) 0.1% FA in water; (B) 80% ACN, 0.1% FA in water. The peptides were eluted directly into the Q-exactive HF-X mass spectrometer that was operated in data-dependent acquisition mode with selection of Top-N 20 most abundant peptides. The parameters used were as following. Peak width 15s, use lock masses was set to off. The full MS scans were performed at 120000 resolution (FWHM at 200 m/z), AGC target 3e6, maximum injection time 100ms, scan range 400 to 1210 m/z, spectrum data type: Profile. For ddMS2, the resolution was 15000 FWHM at 200 m/z, AGC target was 1e5, maximum ion injection time (IT) was 25 ms, loop count was 20, MSMS scan range was 200 to 2000, normalized collision energy (NCE) was set to 28.

For the cancer cell SRL, the high pH RP purified Cys-peptide fraction were each resolubilized in 0.1% formic acid and a fraction was loaded onto a disposable stage-tip pre-column according to the manufacturer’s protocol (Evosep, Odense, Denmark). Peptides were separated on an Evosep analytical column using a 21 minutes generic method provided with the Evosep instrument. Peptides eluted using the 21 min gradient from the stage tip pre-column were analyzed on a Q-Exactive HFX (Thermo Fisher Scientific, Bremen, Germany) with the following parameters: MS and MSMS: resolution at 60000 and 7500, AGC target 3e6 and 1e5. Top 40 most intense precursor ions were fragmented after maximum IT of 20 ms, with NCE 28, at an intensity threshold of 1e5.

#### DIA runs

For MS1 based high resolution mass spectrometry data-independent acquisition (HRMS1-DIA) analysis, full MS scan was performed at FWHM 120000 at 200 m/z, MS AGC target of 3e6, maximum IT 50 ms, scan range 400 to 1210, spectrum type profile. For DIA MS2, default change state was set to 3, resolution 30000 FWHM, AGC target value of 1e6, loop count 18, isolation window 15 m/z, fixed first mass 200 m/z, NCE 28, at an intensity threshold of 1e6, spectrum data type was set as profile.

For the HeLa cell evaluation of the Cys-DIA method normal peptides and Cys-peptides the EASY-LC system, with an EASY-SPRAY 15 cm, 150 ID column, was used with 3 different chromatographic gradients (60, 20 and 10 min).

For the cancer cell comparison, the purified Cys-peptides were analyzed using the HRMS1-DIA method on an Evosep LC system with a 21 min fixed gradient using a stage tip as pre-column. The various Cys-peptide samples were loaded onto a Stage tip pre-column according to the manufactures protocol (Evosep, Odense, Denmark). The Cys-peptides were eluted from the stage tip onto an analytical column directly into a Q-exactive HFX MS system (Thermo Fisher Scientific, Bremen, Germany). The Q-exactive HFX was operated as described for the HeLa peptides DIA above.

### Data analysis

#### Spectral libraries generation

To generate the sample specific spectral libraries, DDA raw files from the high pH RP fractionated samples were searched in Proteome Discoverer (PD) 2.3 (Thermo Fischer Scientific) using Mascot and Sequest HT search algorithms. The raw files were searched against a Human database from SwissProt (May 2019). Trypsin was used as proteolytic enzyme with a maximum of 2 missed cleavages, a precursor mass tolerance of 10ppm, and fragment mass tolerance of 0.02Da, minimum peptide length of 6 and maximum peptides length was 150 amino acids. Oxidation (M) and acetylation (Protein N-term) were selected as dynamic modifications whereas Carbamidomethyl(C) for Cysteine was selected as a static modification during the analysis. The Percolator algorithm^17^ was used as a filter for the high confident identification with 1% false discovery rate (FDR).

The resulted pdResult files from the PD 2.3 searches were uploaded to the Spectronaut X™ software (Biognosis) with option to generate library from Proteome Discoverer according to the instructions given in the User Manual of Spectronaut X™. The raw files were assigned and the FASTA file for Human was downloaded from SwissProt in May, 2019 and a FASTA file for iRT was selected for searches. The libraries were generated with default BGS Factory settings with the exception for the cysteine specific libraries where in Spectral Library Filters only the peptides with Carbamidomethylation on Cysteine (C) as modification were allowed for the spectral library generation. This resulted in the generation of cysteine libraries with 100% cysteine containing peptides.

#### DIA data analysis

The resulting raw files from HRMS1-DIA analysis were loaded into the Spectronaut X™ software. The project specific spectral libraries were assigned to each sample. These spectral libraries provide actual fragment ion intensities and help to identify the peptides with high sensitivity and accuracy. The DIA files were searched with default BGS Factory Settings with the exception that we used MS1 level for quantification. The FASTA files for Human and a FASTA file for iRT were selected for DIA searches as required. Briefly, for data extraction, dynamic mass tolerance strategy at MS1 and MS2 was used with correction factor of 1. The XIC RT Extraction Window was set to dynamic with 1 as correction factor. For Calibration, the calibration mode was set to Automatic. The source specific iRT Calibration and Used Biognosis iRT Kit were enabled. For identification, Scrambled and Dynamic were used as decoy methods. Kernel Density Estimator was applied as a *p* value estimator. Major and minor grouping was by Protein Group ID and Stripped Sequence respectively. The major group quantity was set to mean peptide quantity whereas for minor group quantity mean precursor quantity was used. MS1 level was used for quantification with Area as Quantity Type and q-value was used for data filtering. The Cross Run Normalization was enabled using Local Normalization as normalization strategy with q-value sparse for Row Selection. The Protein Inference Workflow was set to Automatic.

### Statistical analysis and data interpretation

After the DIA searches complete in Spectronaut, the data were exported at the protein or peptide level for further analysis. The cancer cell line data was uploaded into Perseus^18^ and log2 transformed. Furthermore, different scatter plots and histograms were made to see the data quality. For statistical analysis of the cancer samples, the files with log2 transformed protein intensities were uploaded to an in-house shinyApp for Limma and RankProduct test^9^. A p-value cut-off of 0.01 and fold change of 2 was used for the statistically significant regulation. The resulting lists of regulated proteins were analyzed in MetaCore for pathways and disease biomarker networks for the cancer samples. WikiPathways analysis was also used for the significantly regulated proteins in M4A4 and CL16 cancer cell lines compared to the NM2C5, using CluePedia and ClueGO as a Cytoscape plug-ins^19, 20^.

## Acknowledgements

Thermo Fischer Scientific is acknowledged for a Postdoctoral Fellowship. This study was supported by the Villum Center for Bioanalytical Sciences at SDU.

